# Defecation in preparation for ecdysis drives microplastic clearance in cricket nymphs

**DOI:** 10.64898/2026.01.08.698425

**Authors:** Jennie E. Mills, Marshall W. Ritchie, Émile Vadboncoeur, Heath A. MacMillan

**Affiliations:** Department of Biology, Carleton University, Ottawa, ON, Canada, K1S 5B6

**Keywords:** pollution, microplastic, moulting, Orthoptera, development, ecdysis

## Abstract

Plastic pollution is widespread in both terrestrial and marine environments, creating significant ecological concerns. Animals that occupy lower trophic levels, like many small insects, can ingest and retain plastics for extended periods before eliminating them. Ecdysis, or moulting, occurs in arthropods during development and facilitates growth, but its role in microplastic (MP) clearance, and whether it is impacted by plastics, are largely unexplored. We used the cricket *Gryllodes sigillatus* to examine how moulting influences MP clearance in a hemimetabolous species known to ingest and tolerate MPs throughout development. We tested two competing hypotheses of how moulting influences plastic clearance: (1) that moulting of the gut lining removes MPs, or (2) that cessation of feeding and purging frass before ecdysis occurs removes MPs. In doing so we also provide new evidence that cricket nymphs exhibit a cyclical (cosinor) pattern of frass production associated with ecdysis. MPs were eliminated from the cricket’s gut before ecdysis, driven by the cessation of feeding and cyclical frass production that clears the gut in preparation for ecdysis. We also observed that the timing of ecdysis and overall frass production remained unaffected by continuous MP ingestion, whereas switching between MP and non-MP diets caused modest changes in frass timing within an instar. Crickets are known to biofragment MPs into NPs, which, with gut-clearing, likely allows them to deposit plastic-laden frass rapidly and reduces the likelihood of upward trophic transfer by reducing plastic retention.

## 1. Introduction

Plastics are artificially synthesized polymers widely used in the food, chemical, transportation, and construction industries (Andrady and Neal, 2009; Saleh, 2021; Plastics Europe, 2024). Polyethylene (PE), polystyrene (PS) and polyvinyl chloride (PVC) are among the most prevalent polymers synthesized (Geyer, Jambeck and Law, 2017; Plastics Europe, 2024). Globally, plastic production and consumption have been rising, from approximately 2 megatons in the 1950s to 400 megatons in 2022 (Houssini, Li and Tan, 2025). Unfortunately, widespread mismanagement and inefficient recycling of plastics are expected to produce ∼12,000 Mt of discarded plastic waste by 2050 (Geyer, Jambeck and Law, 2017). Over time (estimated at hundreds of years; An et al., 2020; Barnes et al., 2009), these discarded plastics break down through natural biotic and abiotic processes (Cai *et al*., 2023; Emenike *et al*., 2023; Song, Wang and Li, 2024). This process contributes to the production of microplastics (MPs; 5 mm to 1 µm) and nanoplastics (NPs; 1 µm to 1 nm) (Peng *et al*., 2023), which persist in the environment.

Small animals, like arthropods, living in contaminated environments regularly ingest MPs and NPs (Devriese *et al*., 2015; Welden and Cowie, 2016; Abbasi *et al*., 2018; Walkinshaw *et al*., 2020). The retention time of these plastics in the digestive system varies widely across animal taxa and likely depends on particle size (Rist, Baun and Hartmann, 2017; Kang *et al*., 2021; Yu, Nakaoka and Chan, 2021). Within Arthropoda, mealworms retain ingested 150-300 µm MP particles for up to 48 h (Peng et al., 2023; Peng and Wang, 2024), while shore crabs can retain 10 µm MP particles for up to 14 days (Watts *et al*., 2014). As they continue to ingest plastics, these small animals can accumulate concentrations that are proportionally higher than those of organisms at higher trophic levels, up to 16 times higher per unit body mass (Walkinshaw *et al*., 2020). However, the level of accumulation also depends on the animal’s plastic retention time, which influences the rate at which accumulation is offset by excretion of plastic particles back into the environment (Watts *et al*., 2014; Krais *et al*., 2022; Fritz *et al*., 2023; Peng *et al*., 2023; Peng and Wang, 2024). Such differences in retention times also have important implications for plastic bioavailability in ecosystems and can mitigate or facilitate trophic transfer. Thus, it is important that we clearly understand what factors, such as the life history traits of a species of interest, influence plastic retention.

A largely unexplored factor that could affect the retention time of MPs in arthropods is ecdysis (Allison *et al*., 2025) – the process by which most arthropods moult/shed their exoskeleton during larval or nymphal growth (Fahrbach, 2010). The Norway lobster retains MPs until it sheds its gut lining during ecdysis, at which time the plastic particles are cleared from the gut (Welden and Cowie, 2016). Most insects are also known to shed two regions of their gut lining (exoskeleton) during gut epithelium moulting: the foregut and hindgut (Engel and Moran, 2013; Hammer and Moran, 2019). There is also an acellular peritrophic membrane that protects the midgut epithelium of most insects from abrasive gut contents, digestive enzymes, and pathogens (Berenbaum and Isman, 1989; Erlandson, Toprak and Hegedus, 2019; Nunes, Sucena and Koyama, 2021) and this lining is also shed regularly during ecdysis (Berenbaum and Isman, 1989; Engel and Moran, 2013; Nunes, Sucena and Koyama, 2021). At this time, the peritrophic matrix and inner epithelial cell layer are degraded and replaced (Berenbaum and Isman, 1989; Nunes, Sucena and Koyama, 2021). These events occur during every moult in most arthropods, including the Norway lobster, so shedding of the gut epithelium may broadly impact the fate of ingested MPs in arthropods.

In preparation for ecdysis, arthropods undergo behavioural changes, such as ceasing food consumption, which permits gut voiding (Ayali, 2009). In holometabolous (complete metamorphosis) insects like the tobacco hornworm (*Manduca sexta*), the increase in ecdysone at the end of the last larval instar initiates a fasting period and purging of gut contents hours to days before metamorphosis to the adult life stage begins (Nijhout, Davidowitz and Roff, 2006; Suzuki *et al*., 2013). Ecdysone is a steroid hormone that regulates ecdysis in insects (Cabej, 2012). Shifts in feeding and frass production could affect the retention time of plastics and other pollutants during development, as they occur at each ecdysis event. This pattern of surges in ecdysone, ceased feeding and gut purging is best studied in holometabolous models, but has also been documented in hemimetabolous (incomplete metamorphosis) Orthopterans (lubber grasshopper; *Romalea microptera)* before moulting in every instar (Rackauskas, 2006; Riddiford, 2012). This purging of the gut during ecdysis could therefore naturally clear contaminants like MPs or NPs from the digestive tract before the gut exoskeleton is shed.

Crickets (Orthoptera) are an excellent model organism for studying how ecdysis influences plastic retention. They are a diverse group of terrestrial arthropods and an abundant prey item (Knowlton and Harmston, 1943; Zuk and Kolluru, 1998; Pinto and Torres-Carvajal, 2023) with global distributions (Magara *et al*., 2021). Crickets are hemimetabolous insects that develop through a series of nymphal instars before reaching adulthood and are common model systems in the study of development and behaviour. Each instar lasts several days and ends with a moult triggered by surges in ecdysteroid hormones (Riddiford, 2012). Cumulative processes across nymphal instars allow progressive mass gain and the eventual adult emergence (Clifford, Roe and Woodring, 1977; Woodring, Roe and Clifford, 1977; Woodring, Clifford and Beckman, 1979; Roe, Clifford and Woodring, 1980; Kong *et al*., 2025). Within each nymphal instar, cyclical physiological changes occur that enable crickets to acquire, process, and allocate resources for growth and eventual ecdysis. During the first half of an instar, food consumption and digestive activity peak, leading to maximal growth and high mass output (Woodring, Roe and Clifford, 1977; Woodring, Clifford and Beckman, 1979). For example, mass in the penultimate instar of the cricket (*Acheta domesticus*) can increase by approximately 150% within the first four days (Woodring, Roe and Clifford, 1977). By contrast, feeding and frass production decline sharply in the latter half of an instar as ecdysis approaches. By the end of the instar, crickets cease feeding and excreting but remain metabolically active, relying on stored energy reserves and gradually losing body mass until ecdysis is complete and the cycle begins anew (Woodring, Roe and Clifford, 1977; Woodring, Clifford and Beckman, 1979).

The tropical house cricket (*Gryllodes sigillatus*) tolerates ingestion of MP beads as early as two weeks after hatching and grows to a similar adult body size on diets containing as much as 10% w/w PE (Fudlosid *et al*., 2022; Ritchie, 2024). Between three and four weeks of age (at 30-34°C), they undergo rapid growth, progressing through instars 5-7 before reaching their adult instar (instar 8) around 5-6 weeks of age (Kong *et al*., 2025). This period of frequent moulting provides an opportunity to use this model species to better understand the role of ecdysis in plastic clearance under controlled conditions.

We first observed that nymphs contained less plastic in their digestive systems during ecdysis, but this clearance occurred before exoskeletal moulting (Figure S1). We therefore aimed to directly examine the relationship between ecdysis and plastic retention by tracking the presence of plastic in nymphs throughout an entire instar. We tested two competing hypotheses: (1) that clearance of the gut exoskeleton clears MPs, or 2) that ceasing feeding and clearing frass before ecdysis clears MPs. We predicted that nymphs would increase frass production immediately before moulting, and that PE plastics would be detected in pre-moult frass but not in the post-moult frass or digestive tracts unless crickets were allowed to ingest MPs after ecdysis occurred.

## 2. Methods

### 2.1 Care and Feeding

Our colony of *G. sigillatus* was established from eggs sourced from Entomo Farms (a commercial producer of crickets as food and feed). The colony is reared in bins (60 cm x 40 cm x 31 cm) held at a density of ∼500 crickets and maintained at ∼30 ± 2°C and 40% RH with a 14:10 L:D light cycle at Carleton University (Ottawa, ON). Egg cartons were provided for shelter for the duration of cricket development. In the colony, crickets were given water and fed *ad libitum* with a standard feed of ground wheat, corn, soymeal, and oat hulls (16% Layer Pellet, Ritchie Feed and Seed Inc, Ottawa, ON). The same diet was used as the control diet in experiments, and the MP diet was the same basic diet, but contained 1% w/w low-density ∼100 μm diameter fluorescent blue polyethylene microspheres (Cospheric, Fluorescent Blue Polyethylene Microspheres 1.13 g/cc 90–106 μm).

### 2.2 Effects of moulting on microplastic retention in the cricket nymph gut

#### 2.2.1 Moult -tracking experimental setup

To confirm whether ecdysis aids in plastic clearing, we placed 30 nymphs (2-3 weeks) each in four plastic bins (two for control and two with plastic feed) and allowed them to feed for 24 h. Immediately after the 24 h feeding period (D0), we randomly selected 10 individuals from each bin and dissected out their digestive tracts as described in Ritchie *et al*. (2023). Briefly, for dissections, we anesthetized crickets by exposing them to CO_2_ for 10 seconds, washing them in a dish of water, and placing them in saline solution. We then dissected the crickets under a dissecting microscope (Stemi 508, Zeiss, Oberkochen, Germany) by pinching the crop and gently teasing out the digestive tract. Finally, we carefully removed the gut with forceps, retaining its contents, and placed it in a 1.6 ml microcentrifuge tube that was stored at -20°C until further analysis. All remaining individual nymphs were then transferred to 3-oz deli cups and held in an incubator (32°C, 40% RH, 14:10 L:D light cycle) for the remainder of the experiment, so that we could track individual moulting events. Individuals were given *ad libitum* access to water and control feed and were checked 1-2 times daily to track moulting until ∼50% of individuals had moulted (∼2-3 d). This design allowed us to compare microplastic loads in crickets that had undergone ecdysis and those that had not, while controlling for time. We identified moulting events by the presence of whole or partial moulted exoskeletons within the cups, as many nymphs began to consume their moults shortly after shedding, leaving small pieces of the moult behind in the cups. When a moulting event was confirmed, we performed gut dissections immediately as described above. We then repeated these methods for a second age-group of nymphs (3-4 weeks). One nymph from the 2-3-week age-group died and was removed from future analyses. In general, crickets tolerated plastic feeding very well and survival remained high throughout the experiment, as we have noted previously in this species (Fudlosid *et al*., 2022).

#### 2.2.2 Tissue Digestions and MP Imaging

We digested tissues and imaged microplastics contained within them according to the methods of Ritchie et al. (2023). Briefly, we thawed gut samples to room temperature and digested them in either 300 µL (2-3 week-old crickets) or 400 µL (3-4 week-old crickets) of 10% KOH at 60°C for 2 d. After the incubation period, we vortexed each sample and filtered it through 1 µm pore-size glass fiber filters (APFB04700, MilliporeSigma Canada) over a manual vacuum pump set-up. Then we transferred each filter to a clean petri dish and photographed it under a dissecting microscope (Stemi 508 equipped with an Axiocam 105 colour, Zeiss, Oberkochen, Germany) through a 450-nm long-pass emission filter with a 400–415-nm excitation light source (Stereo Microscope Fluorescence Adapter, Nightsea LLC, Lexington, United States). Each resulting image was then examined to confirm the presence of MP particles.

### 2.3 Gut purging in preparation for moulting as a mechanism of microplastic clearance

#### 2.3.1 Frass collection and moult-tracking experimental setup

The results from our first moult-tracking experiment (2.2) indicated that while ecdysis probably contributed to the removal of MPs from nymphs, this removal occurred before they shed their exoskeleton. This observation led us to explore two alternative mechanisms behind this removal: increased frass production approaching ecdysis or the shedding of the gut exoskeleton during ecdysis. We pseudorandomly selected 60 3-week-old nymphs from the colony and identified them as 4^th^ instar based on age and the lack of visible ovipositors in any individuals (Kong *et al*., 2025). Employing a technique from (Jemec Kokalj, Nagode, *et al*., 2024), we immobilized the nymphs with a small piece of cheesecloth and marked the dorsal side of their thorax with a black permanent marker through the material. This mark facilitated the identification of moulted individuals in a group rearing setting throughout the experiment, as the ink would be shed with the exoskeleton (Figure S2). The crickets were divided evenly into two groups: one group was fed a control diet, and the other was fed a 1% w/w plastic-contaminated diet (3 replicate bins of 10 individuals per group). Bins were held in an incubator (32°C, 40% RH, 14:10 L:D light cycle) for daily monitoring.

We monitored the bins in brief intervals (5-10 min) four times per day (∼00:00h, 06:00h, 12:00h, 18:00h) to identify moulted individuals, identified by the absence of a black marker on the thorax. The day and time of the first moulting event into the 5^th^ instar (M1) were recorded for each cricket. Newly 5^th^ instar individuals were then marked again on the dorsal side of the thorax and placed into individual 3-oz deli cups to enable tracking of individual cricket frass production and their next moulting event into the 6^th^ instar (M2) in relation to M1. We defined 5^th^ and 6^th^ instars using a published visual representation, but did not account for sex in this experiment because it is not identifiable in this species until later instars (Kong *et al*., 2025).

Each cup was modified to have a 4 mm x 4 mm mesh wire bottom and was nested within a second “collecting” cup, such that frass fell through the mesh into the second cup, facilitating frass collection (Figure S3). The feed was placed in a raised cap to limit the cricket’s ability to walk in the feed and to prevent MPs from adhering to its exoskeleton. Each cup also contained a piece of egg carton for shelter, and *ad libitum* food and water, and they were held in an incubator (32°C, 40% RH, 14:10 L:D light cycle).

#### 2.3.2 Post-Moult Feeding

After M1 and isolation of crickets into individual cups, the experiment proceeded in a fully crossed feeding design (Fig. 1) to allow us to evaluate any presence after ecdysis M1 (MP-Control) and the effects of a switched diet (MP-Control, Control-MP). Post-M1, we kept half of the individuals from each initial diet group on the same diet (Control – Control, MP – MP), and the other half were switched to the opposite diet (Control – MP, MP – Control), ensuring that the final treatment groups included individuals from each of the replicate bins. This resulted in four treatment groups, each with 15 individuals, tracked as they progressed toward M2.

**Figure 1.**
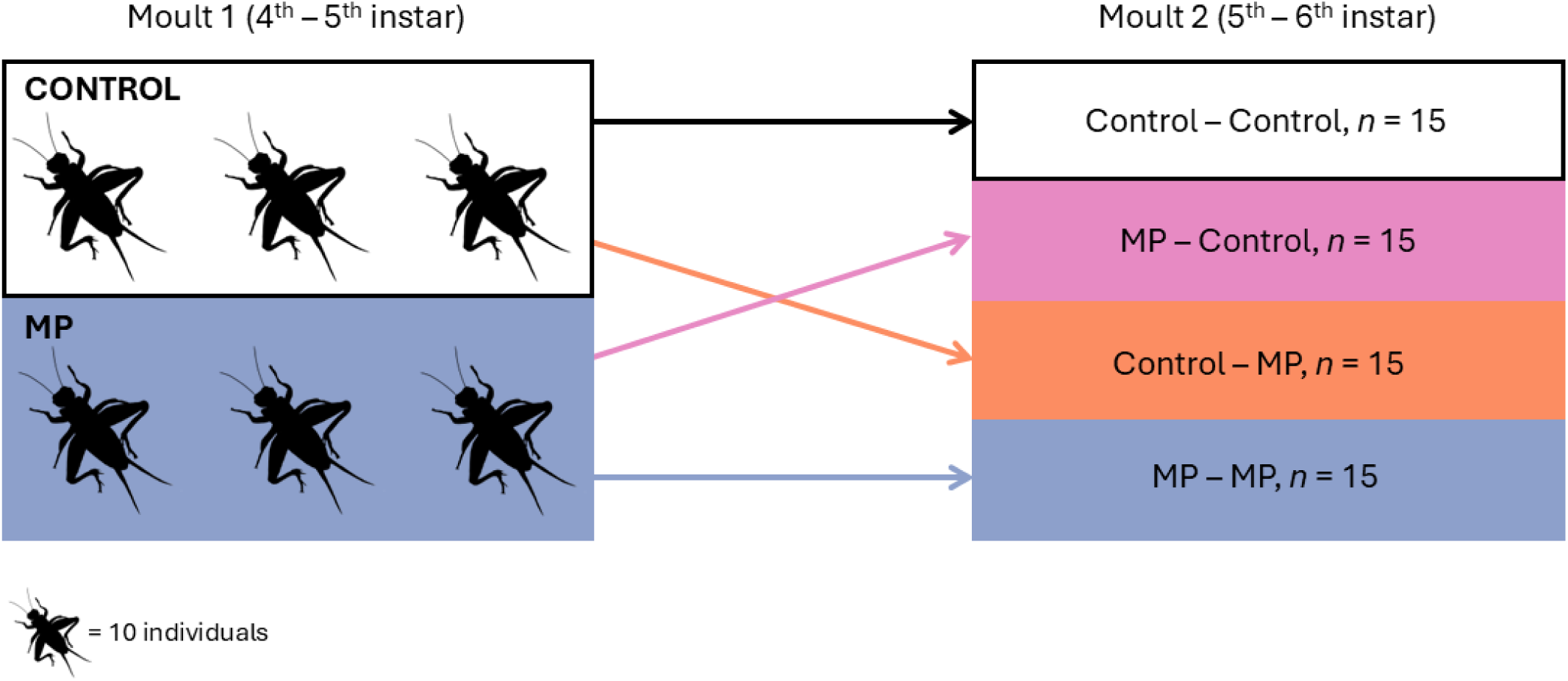
Fully factorial experimental feeding design to test how moulting events in *G. sigillatus* contribute to plastic clearance when fed a consistent or switched diet.

#### 2.3.3 Individual Monitoring and Frass Collection/Processing

We inspected each cricket every 6 h starting after M1 (∼00:00 h, 06:00 h, 12:00 h, 18:00 h). During each inspection, we gently shook the cups, allowing any stuck frass to dislodge and fall through the mesh into the collecting cup. Then we separated the mesh-bottomed inserts from the collecting cup, collected the accumulated frass and transferred it to pre-dried and pre-weighed 1.6 mL microcentrifuge tubes. Then we placed the tubes in a drying oven at 60°C for 24 h. Next, we weighed the samples within the tubes (Sartorius Entris II Analytical Balance, BCE124I-1S) to 0.001 g and quantified the frass dry mass by subtracting the pre-measured tube mass from the final mass. All frass was frozen at -20°C for future analysis. One control cricket died 18 h after M1; its data were excluded from all analyses.

We conducted frass collection and moulting checks at the same frequency until all individuals had moulted a second time (M2), and the day and time of each moult were recorded. We kept the crickets in their cups for an additional 6 h after M2 to test for MPs in the frass immediately after a moult. We euthanized each cricket by freezing and stored them at -20°C for future analysis of the gut content of MPs.

#### 2.3.4 Frass and Tissue Digestions, MP Filtration and Image Analysis

All frass collected during the experiment was stored for later analysis. During the experiment, we observed that the nymphs ceased frass production before and during moulting, as is seen in other Orthopterans (Rackauskas *et al*., 2006). Thus we chose to analyze the MP content of the first frass sample after M1, the sample at 48 h post-M1 (average peak frass production), the last sample before M2, and, if present, the sample after M2 for all individuals. We analyzed the plastic content in the frass following published methods (Ritchie et al., 2023). Briefly, we thawed the frass samples to room temperature, isolated one frass pellet from each (most had only one) and transferred it to a 0.6 mL microcentrifuge tube. Next, we added 30 µL of 10% KOH to each tube, and digested the frass for 48 h at 60°C. Finally, we thawed frozen crickets to room temperature, dissected out the guts and digested them.

All digested frass and gut samples were filtered and imaged following the same procedure outlined in 2.2.2. These images were used to determine whether plastics were present within the individuals’ frass and digestive tracts.

### 2.4 Statistical Analyses

#### 2.4.1 Analyzing the effects of moulting on plastic presence

Differences in the proportion of particle-positive individuals between moulted and non-moulted groups were evaluated by comparing plastic presence in nymphs directly to the baseline day (D0). Individuals were classified as D0, moulted, or non-moulted, and overall group differences were assessed using binomial GLMs of the presence of plastic and the effects of moulting for both the 2-week and 3-week data. Overall moulting time was evaluated using a GLM and a type III ANOVA. All statistical analyses and plotting were done in R v4.4.1(R Core Team, 2024).

#### 2.4.2 Cosinor curve

To evaluate whether frass production exhibited rhythmic patterns and whether these rhythms differed among diets, we used a mixed-effects cosinor model. Time (hours (h) since the start of sampling) was transformed into cosine and sine components using a 124-hour cycle (the period of the dataset determined using the function nlsLM from the R package minpack.lm) to represent the oscillation’s amplitude and phase. These variables allow linear mixed models to quantify both the amplitude and timing (phase) of rhythmic behaviour. We then fit a mixed-effects model of the form: frass ∼ diet × (cosine + sine h) + (1 | ID). Here, diet and its interactions with the rhythmic terms were tested to determine whether treatments altered the amplitude or timing of the frass production cycle. Individual cricket nymph IDs were included as a random effect to account for repeated sampling. Model fitting was performed using Satterthwaite-adjusted t-tests (lmerTest) and Type III Wald χ² tests used to assess statistical significance (Hou, Tomalin and Suárez-Fariñas, 2021).

To quantify rhythmicity within each diet group, we calculated R² values from separate linear cosinor fits using a fixed 124-hour cycle. Parameters of the cosinor model were calculated in R from published methods (Refinetti, Cornélissen and Halberg, 2007; Hou, Tomalin and Suárez-Fariñas, 2021) and equations used are presented within the supplemental material. All analyses were conducted in R (version 4.5.1) using the lme4, lmerTest, and car packages.

## 3. Results

### 3.1 MPs are cleared from the gut before exoskeletal moulting occurs

We wanted to investigate the relationship between plastic retention and ecdysis. When cricket nymphs were fed on MP-contaminated diets, most individuals (75-90%) contained detectable plastic particles on Day 0 (active feeding on MP diet), regardless of age. Once ∼50% of crickets had undergone ecdysis, proportions of both moulted and non-moulted individuals containing detectable plastics were significantly lower (13-35%; *χ*²(2) = 20.85, *p* < .001; Table 1). 3-4 week-old moulted individuals tended to contain plastics more than 2-3 week moulted individuals, but this difference was not statistically significant (*z* = 1.61, *p* = 0.107; Table 1). At 2-3 weeks of age, plastic content in the digestive system was lower than on Day 0, regardless of whether the crickets had moulted (*z* = –3.49, *p* < 0.001) or not (*z* = –3.41, *p* < 0.001). Similarly, 3-4 week-old crickets were less likely to contain plastics in their digestive system, whether they moulted (*z* = – 3.11, *p* = 0.0019) or not (*z* = –4.23, *p* < 0.0001). The number of particles present in crickets at the end of the experiment was also notably low (N ≤ 8; Figure S1). No plastics were found in crickets fed the control diet. When intact moults were recovered from MP-fed crickets, plastic particles were often adhered to the moult, likely due to contact with contaminated food. This would provide individuals with the opportunity to re-ingest MPs when consuming their shed exoskeleton. We therefore expected that this might account for the presence of plastic after a moult. It is also possible that the low counts observed in non-moulted individuals resulted from reingestion of MPs during grooming.

**Table 1.** Presence of microplastic (MP) in non-moulted and moulted cricket nymphs (2-3 and 3-4 week *G. sigillatus*). Asterisks (*) represent significant differences in the values compared to the Day 0 group.

| Age | Group | # nymphs containing MP | % nymphs containing MP |
| --- | --- | --- | --- |
| 2-3 weeks | D0 | 15/20 | 75 |
|  | Non-moulted (~D1-3) | 3/19* | 15.7* |
|  | Moulted (~D1-3) | 3/20* | 15* |
| 3-4 weeks | D0 | 18/20 | 90 |
|  | Non-moulted (~D1-2) | 3/23* | 13* |
|  | Moulted (~D1-2) | 6/17* | 35* |

During dissections of moulted 3-4 week-old crickets, we noted that nine individuals were actively shedding their proventriculus and foregut exoskeleton within the digestive tract (Figure 2A, B). In these cases, care was taken to dissect the gut while retaining the orientation of the moult within the gut, and moults were always inverted relative to the typical orientation of the proventriculus and foregut (Figure 2C). This suggests that gut ecdysis begins in the foregut, which is pulled through itself as it passes through the digestive tract for eventual excretion. The presence of the shed gut exoskeleton in the frass supports this, as seen in other individuals at a later stage of ecdysis (Figure 2D, E).

**Figure 2.**
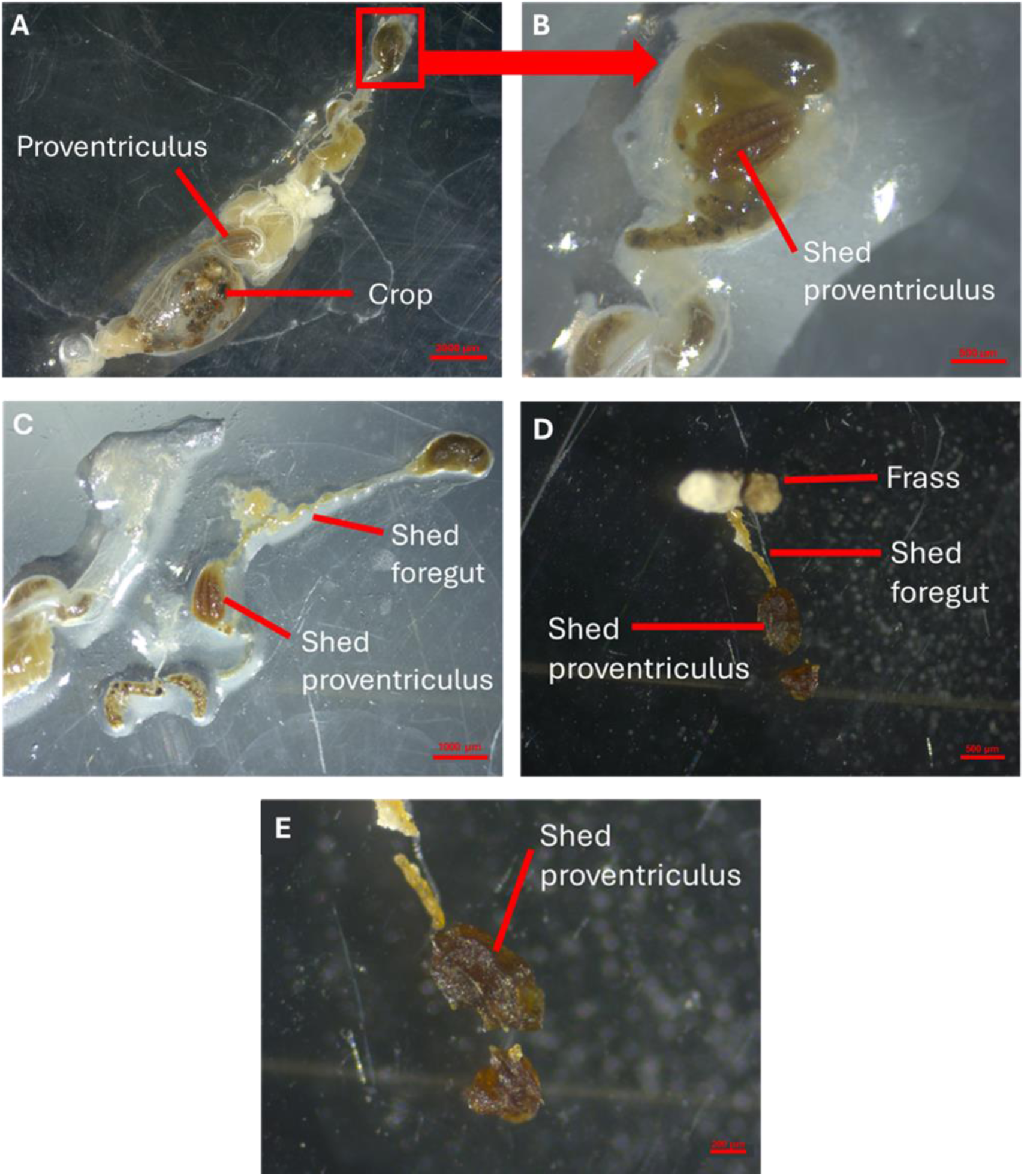
The cricket gut exoskeleton is moulted in addition to the outer exoskeleton and did not contain microplastics when it was shed. (A) Example image of an excised, whole cricket gut from a 3–4-week nymph that had moulted and consumed its shed exoskeleton; red box indicates the portion of the hindgut containing the shed proventriculus and gut exoskeleton. (B) A close-up image of the hindgut clearly showing the presence of the shed proventriculus. (C) Image of the shed proventriculus and foregut exoskeletons dissected from the hindgut; the shed was placed in the same orientation as it was found inside the hindgut. (D) Image of a frass pellet from a 3-4 week nymph adhered to a shed proventriculus and foregut exoskeleton. (E) Close up of the shed proventriculus lining.

In individuals actively shedding the gut exoskeleton, we observed no MPs in gut moults or further down the digestive tract of crickets fed MPs (Figure 2). This implied that MPs were already cleared from the gut before the lining was shed, apart from the consumed exoskeleton, which could have adhered MPs (visually confirmed by the presence of chitinous pieces with setae, located inside the crop). The digestive tracts of most moulted individuals were relatively empty of food or digested material (Figure 2A). Together, these results suggested that gut contents are cleared from cricket nymphs before moulting of the exoskeleton, and that the shedding of at least part of the exoskeleton in the foregut and proventriculus occurs after moulting of the external exoskeleton.

### 3.2 MP feeding does not impact time to moulting

If ingested MPs are cleared before the gut epithelium is moulted, we suspected that they would be removed along with frass before the external exoskeleton is shed. We therefore examined frass production throughout an entire instar and how MP ingestion influenced frass output. Using four treatment groups of cricket nymphs in a fully crossed design (Control–Control, MP–MP, Control–MP, and MP–Control), we tracked frass production throughout the entire fifth instar (Figure 3). There was no significant effect of treatments on time to ecdysis (χ² = .16, *p* = 0.98) between M1 and M2 (Figure 3A). We found that frass production followed a cosine curve and applied a cosinor statistical model (Figure 3B) to the data (Cornelissen, 2014). Cosine curves fit the data well for each diet group: R² = 0.88 for Control, 0.96 for MP, 0.91 for Control–MP, and 0.96 for MP–Control.

**Figure 3.**
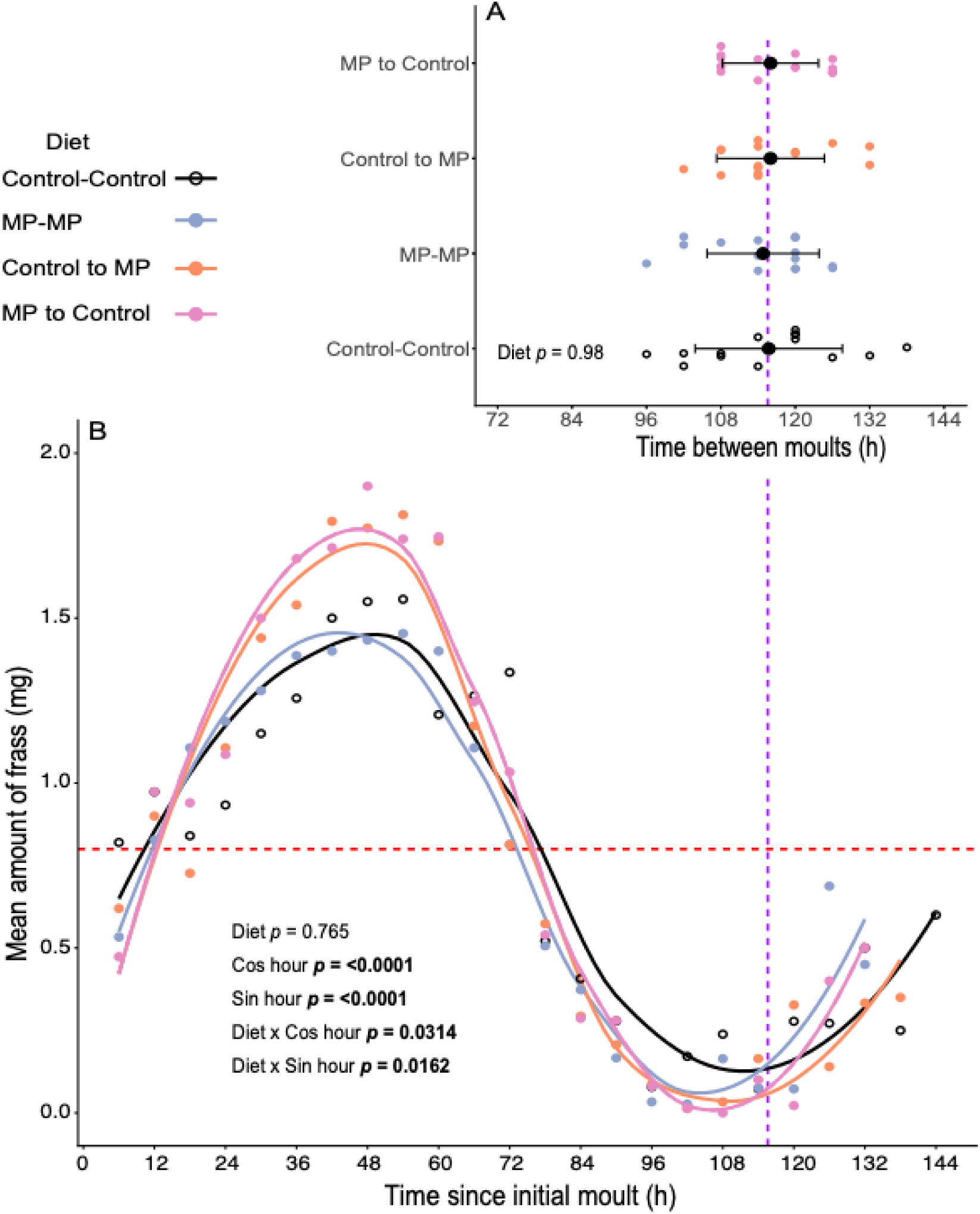
Moulting timing and frass produced by *G. sigillatus* (n= 14-15, per diet) nymphs between instars 5 and 6. (A) Number of hours for individuals (coloured points) to complete instar 5 and moult into instar 6. Solid black points represent the mean ± SE time of moulting on each diet. (B) Mean amount of frass (mg) produced by crickets in each diet (coloured points) with curves of best fit (cosinor) shown for each diet (coloured lines). Dashed horizontal and vertical lines represent the mean MESOR (Midline Estimating Statistic Of Rhythm; red) and the mean time to moult to instar 6 (purple), respectively, which did not significantly differ among diet treatment groups.

Total frass production was not found to be impacted by diets within the model (χ² = 1.15, *p* = 0.77; Table S1). This was found to hold for each diet compared with the control diet for frass production: consistently fed MP-MP (*p* = 0.84) and switched diets Control-MP and MP-Control (*p* = 0.63 and 0.46, respectively; Table S2).

The phase (timing) component of the rhythm (sin hour) was also highly significant (χ² = 129.19, p < 0.0001; Figure 3C), indicating that frass production showed a strong and consistent pattern across the instar. Experimental diet influenced the timing of the oscillation, as there was a significant interaction in the effects of diet and the phase component on frass production (χ² = 10.29, p = 0.016; Figure 3C). Analysis of the fixed-effects model revealed that nymphs on the MP–Control and Control–MP diets exhibited significantly earlier/advanced phases relative to the Control diet (p = 0.0029 and p = 0.0118, respectively; Table 2; Table S2). Although the MP diet showed a numerical phase advance of 2.47 h (Table 2), this shift was not statistically significant (p = 0.0989). Together, these results demonstrate that diet affected the timing of peak frass production, with the most consistent phase advances occurring in groups that experienced diet switches.

**Table 2.** Cosinor-derived rhythmic parameters quantifying the underlying temporal structure of frass production. MESOR represents the rhythm-adjusted mean frass mass, amplitude reflects the oscillation magnitude, and peak and trough times indicate the predicted timing of rhythmic extrema within the 124-hour cycle. Peak and trough values represent the modelled frass mass at these extrema. The peak–trough range summarizes overall rhythmic strength, and phase delay indicates how much earlier each diet reaches its rhythmic peak relative to the Control group. Together, these descriptive and model-derived metrics provide a comprehensive characterization of how diet influences both the magnitude and rhythmic organization of frass production.

| Parameter | Control | MP | Control to MP | MP to Control |
| --- | --- | --- | --- | --- |
| **MESOR (mg)** | 0.77 | 0.75 | 0.83 | 0.85 |
| **Amplitude (mg)** | 0.69 | 0.76 | 0.88 | 0.92 |
| **Peak time (h)** | 45.32 | 42.85 | 44.61 | 44.72 |
| **Trough time (h)** | 107.32 | 104.85 | 106.61 | 106.72 |
| **Peak value (mg)** | 1.47 | 1.51 | 1.70 | 1.77 |
| **Trough value (mg)** | 0.08 | −0.00 | −0.05 | −0.07 |
| **Peak–Trough range (mg)** | 1.39 | 1.51 | 1.75 | 1.84 |
| **Phase advance vs Control (h)** | 0 | 2.47 | 0.71 | 0.60 |

Alongside the observed phase differences, the amplitude component of the rhythm (cos hour) was overall significant (χ² = 96.16, *p* < 0.0001; Figure 3B), indicating frass production showed a clear oscillatory pattern throughout the instar. Diets varied in the strength of their oscillations, and the diet-cos hour interaction significantly affected frass production (χ² = 8.84, *p* = 0.031; Figure 3B). Examination of the fixed-effects model showed that this interaction was driven primarily by the MP–Control group, which exhibited a significantly greater amplitude relative to the Control diet (*p* = 0.0453; Table S2). In contrast, the MP diet (*p* = 0.6142) and the Control–MP diet (*p* = 0.1262) did not differ significantly from Control-Control in amplitude. These results demonstrate that diet affected both when nymphs reached peak frass production (phase differences) and the extent to which frass production varied across the instar (amplitude differences). These differences were mainly driven by the groups in which the crickets received a novel diet was switched at the start of the instar (MP-Control and Control-MP).

### 3.3 Gut contents of crickets are cleared before ecdysis

Frass was examined to detect the presence of plastic at four key locations along the frass production curve. This was 6 h after moulting into instar 5, during peak frass production at approximately 44 h (Table 2), the last frass produced before moulting into instar 6, and 6 h after completing moulting into instar 6 (Figure 4). The two groups fed the MP diet (MP-MP and Control-MP) during the 5^th^ instar had a higher proportion of individuals containing plastics at the beginning of instar 5, indicating that within 6 h of ingesting the MP diet, they excreted the ingested plastics. The MP–Control group (fed plastic during 4^th^ instar) had a single individual (out of 15) with plastic in its frass after ecdysis to the 5^th^ instar (M1), suggesting that the ecdysis process does result in almost complete clearance of MPs from the digestive tract.

**Figure 4.**
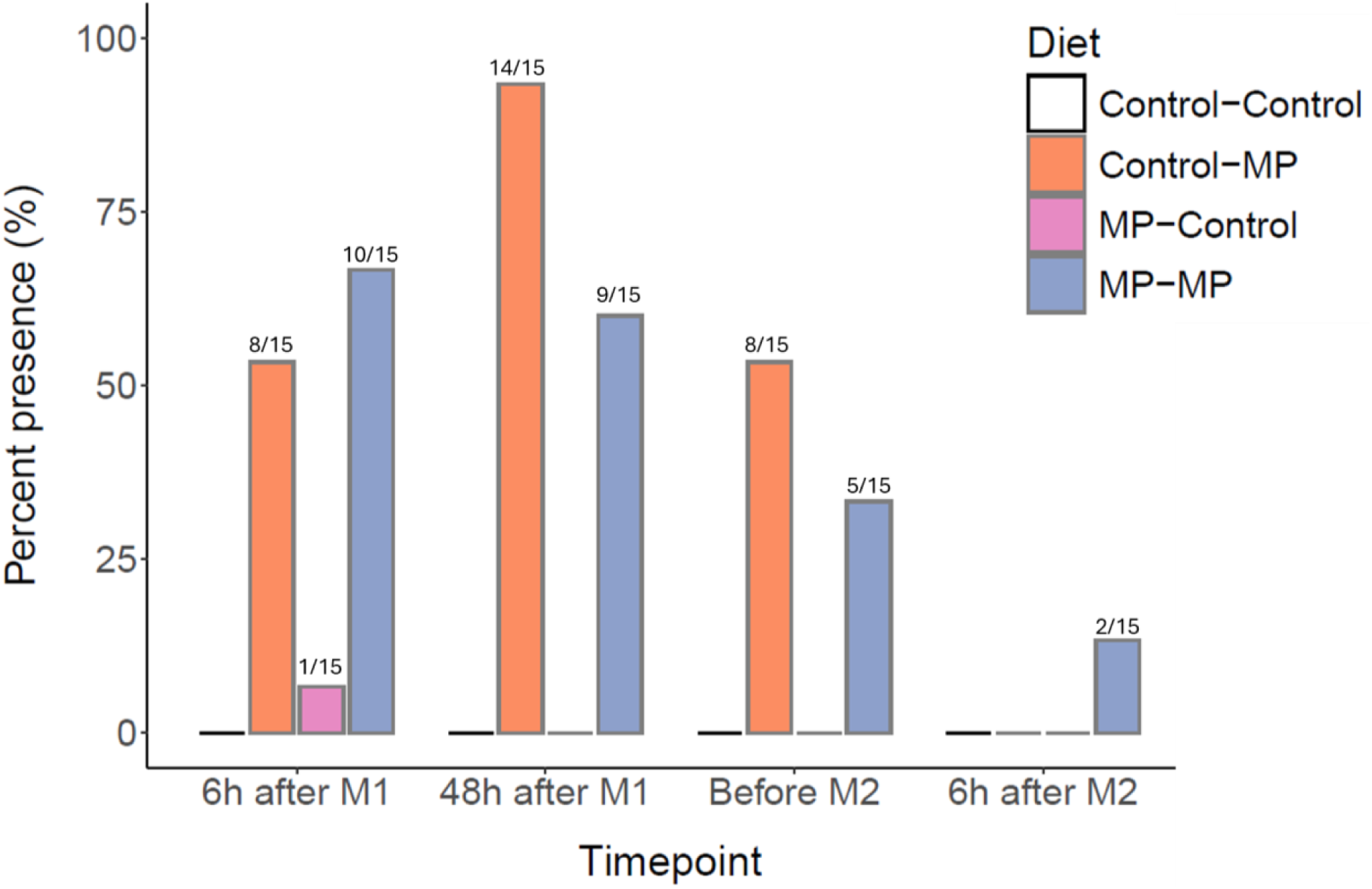
Proportion (%) of frass samples from 3-4 week-old cricket nymphs (*G. sigillatus*) containing MPs. Frass samples were from 6h after moulting event 1 (M1; 4^th^ to 5^th^ instar), the peak average frass production at ∼44 h, the last instance of frass before moulting event 2 (M2; 5^th^ to 6^th^ instar), and in the frass 6 h after M2. Fractions above bars display the number of crickets with positive plastic presence in the frass relative to the total number examined in the group. Groups that do not have a ratio had zero detectable plastics across the entire treatment group.

At the peak frass production (∼44 h), a large amount of plastic was present in the frass of both groups being fed MP diets during instar 5. The Control–Control and MP–Control groups did not contain any plastic at this time, confirming the MPs are not retained after ecdysis to instar 5. After 44 h, frass production dropped sharply, reaching zero (Table S3) just before moulting in every individual (Figure 3). The last frass excreted by crickets in the 5^th^ instar was then examined for plastic content, revealing a significant decrease in overall plastic content between individuals in actively fed MP groups (MP–MP and Control–MP). After examining the frass for plastic content 6 h after crickets successfully moulted to instar 6, only two crickets were found to have excreted plastic: both in the MP-MP group (Figure 4).

## 4. Discussion

In this study, we aimed to understand the effects of the natural progression of ecdysis on the presence of plastics in the guts of cricket nymphs. As previously noted for lobsters (Welden and Cowie, 2016), we found that ecdysis significantly reduced the amount of plastic in cricket digestive systems. Instead of shedding the gut exoskeleton clearing MPs, we found that clearing gut contents before exoskeleton shedding removes most MPs from the digestive tract. We also found that ingesting polyethylene did not affect the timing of ecdysis or total frass production in a cricket.

Other insect nymphs (*Plautia stali)* appear to have difficulty clearing ingested plastic (Oishi, Moriyama, and Fukatsu, 2023). We therefore expected that gut bioaccumulation of plastic could alter the timing of moulting and frass production in a cricket. We instead found that neither process was significantly affected in crickets. A potential reason why the timing of these events is unaffected in cricket nymphs is their ability to biodegrade plastics; crickets biofragment the plastics they ingest, likely through the actions of their mouthparts and proventriculus (Ritchie *et al*., 2024, 2025). This process creates smaller plastics, which are suggested to have longer retention times, but may decrease bioaccumulation by facilitating movement of plastics through the digestive system (Rist, Baun and Hartmann, 2017; Kang *et al*., 2021; Yu, Nakaoka and Chan, 2021).

Feeding on a plastic diet before or during an instar did not affect the timing of ecdysis or total frass produced within an instar of cricket nymphs. We predicted that nymphs would cease feeding and purge their gut contents before moulting, thereby assisting in clearing MPs from the gut. Peak cricket frass production across all groups occurred ∼44 h after ecdysis and following this peak, we found a gradual decrease in frass production for an additional ∼62 h until crickets generally stopped producing frass entirely (Figure 3). This period of low frass production continued until the crickets had successfully moulted to the next instar. These findings match those in *A. domesticus* regarding food consumption over an instar, with peak food intake occurring approximately 48 h after ecdysis and a subsequent decrease over multiple days that ends with zero feed consumption (Woodring, Roe and Clifford, 1977; Woodring, Clifford and Beckman, 1979). The process is much longer in 5^th^ instar *M.* sexta larvae, which cease feeding over an 8 h period and purge their gut contents ∼4 days (96 h) after ecdysis to the 5^th^ larval instar (Dominick and Truman, 1984; Nijhout, Davidowitz and Roff, 2006). This is followed by a 10-30 h wandering period before the larva pupates (Dominick and Truman, 1984; Nijhout, Davidowitz and Roff, 2006), which lasts for 2-3 weeks before eclosion (Dominick and Truman, 1984). Although knowledge on this topic is limited, these findings suggest that differences in the timing of ecdysis-related events and instar timing will impact the ways in which plastics are retained or cleared in different insect taxa.

Switching between MP and control diets in crickets (MP-Control and Control-MP) altered the amplitude and timing of frass production during the 5th instar. This suggests that changes in the available diet, rather than MPs, adjust the timing and peaks of frass production. Switching diets was found to cause a higher peak frass production amplitude but an extended period of no frass production immediately before ecdysis (Figure 3). This difference could be driven by attraction to a novel diet that accelerated feeding early in the instar (Bernays and Bright, 1993). Overall timing of the instar was not affected, nor was the total amount of frass produced within the instar. These observations are also supported in other insects, where ecdysis is not affected at environmentally relevant concentrations in holometabolous insects, such as mealworms (Jemec Kokalj, Dolar, *et al*., 2024), suggesting this highly conserved trait isn’t affected by MP ingestion. Both crickets and mealworms can biofragment MPs (Peng *et al*., 2024; Ritchie *et al*., 2024), which supports prior evidence that crickets don’t experience developmental delays from ingesting plastics (Fudlosid *et al*., 2022; Ritchie *et al*., 2025).

While the ability to clear plastics during ecdysis has been noted in another arthropod (Welden and Cowie, 2016), when during this process plastics are cleared has not been tested in any ecdysozoan. We first found that time spent progressing toward ecdysis was a better predictor of plastic clearance than moulting of the external exoskeleton (Table 1). We also found shed exoskeletons of proventriculi in the hindgut and frass that lacked microplastics (Figure 2). Together, this evidence suggests that removal of the gut exoskeleton occurs after moulting of the external exoskeleton and after microplastics have already been cleared from the gut. The other chitinous material we found in the frass of recently moulted crickets is likely part of the peritrophic membrane lining the cricket gut. The peritrophic membrane starts at the oesophageal valve, located in the esophagus in orthopterans (Wigglesworth, 1930). Clearance of the peritrophic membrane during each consecutive instar moult is noted to help remove pathogens and other microorganisms from insects (Moll *et al*., 2001; Hammer and Moran, 2019), but, at least in crickets, does not appear to be important to microplastic clearance.

## 5. Conclusion

This work demonstrates that the natural process of ecdysis effectively reduces and removes MPs, and possibly other pollutants, from the insect digestive tract. This finding aligns with previous literature on tolerance to MPs in this cricket species. Switching between control and MP feeds influenced the timing and oscillation of frass production within an instar, but MP feeding did not affect total frass production or the timing of moulting to the next instar. While we focused on a specific period of development in this study. We anticipate that these findings would hold across all instars of this species. Given that crickets can biofragment MPs to the NP level (Ritchie et al., 2024) and that gut clearance occurs with each moult, we expect that repeated rounds of ecdysis help to reduce the probability of trophic transfer of ingested plastics to higher trophic levels by reducing retention time. While this could limit upward transfer of MPs through the food chain, consistent release of plastic-laden frass into the environment might also increase the risk of exposure of other animals to excreted NPs.

## Acknowledgments

We would like to thank Sophie Kasdorf for managing colony care and egg rearing and Matthew Tease for his assistance in labelling hundreds of vials.

## Author Contributions

**Jennie Mills:** Conceptualization, Methodology, Validation, Formal analysis, Investigation, Data curation, Writing - Original Draft, Writing - Review & Editing, Visualization. **Marshall Ritchie:** Conceptualization, Methodology, Validation, Formal analysis, Investigation, Data curation, Writing - Original Draft, Writing - Review & Editing, Visualization. **Émile Vadboncoeur:** Conceptualization, Methodology, Investigation, Writing - Review & Editing, Data curation. **Heath MacMillan:** Conceptualization, Methodology, Writing - Original Draft, Writing - Review & Editing, Visualization, Supervision, Funding acquisition.

## Ethics Statement

None.

