## Supplemental material for "Defecation in preparation for ecdysis drives microplastic clearance in cricket nymphs"

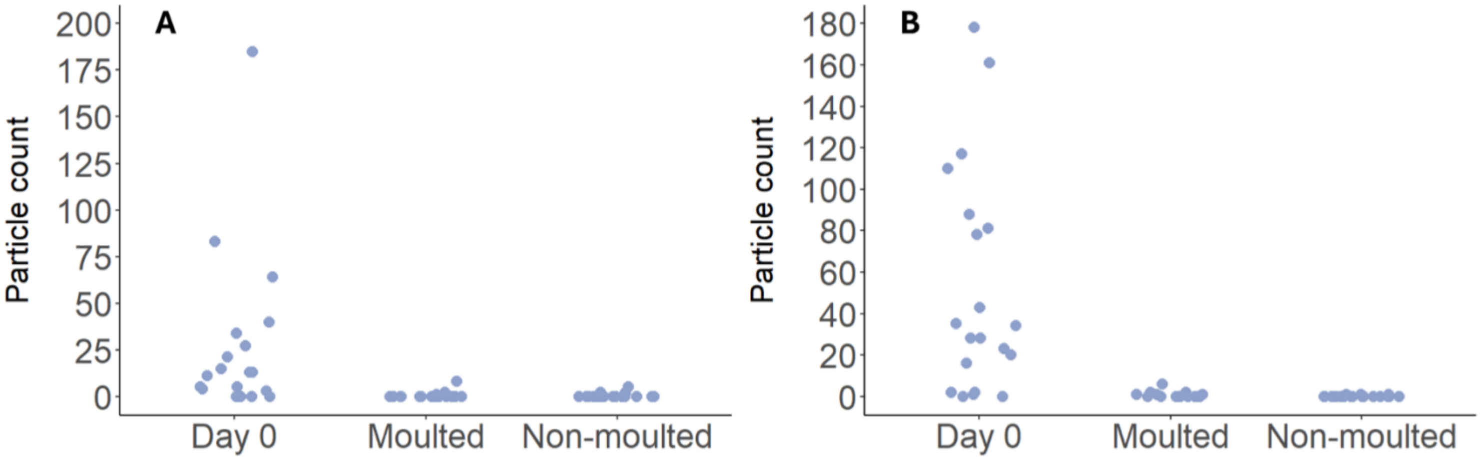


**Figure S1.** Particle counts of MPs within the digestive tracts of nymph crickets between the ages of 2-3 weeks (A) and 3-4 weeks (B). Each point represents an individual cricket. Day 0 is the group sampled at the end of the 24-hour feeding period. Moulted and non-moulted represent when the crickets were sampled after 50% had moulted their external cuticle. This evidence suggested that time spent progressing toward a moult was more important to plastic clearance than moulting itself.

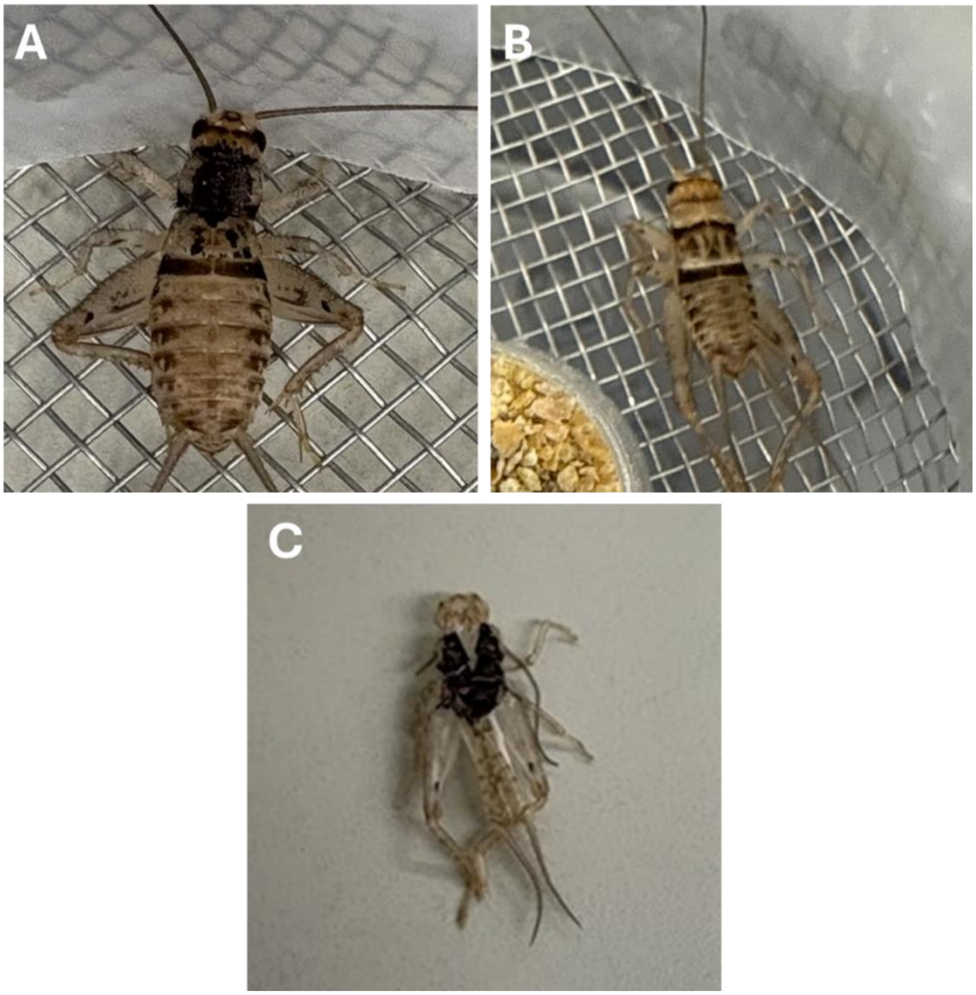


**Figure S2.** (A) A 5$^{th}$ instar nymph with black permanent marker on the dorsal surface of the thorax shortly before moulting to 6$^{th}$ instar. (B) An individual after moulting to 6$^{th}$ instar, showing that the permanent marker ink was no longer present. (C) Intact moulted exoskeleton of another individual showing the permanent marker ink still present.

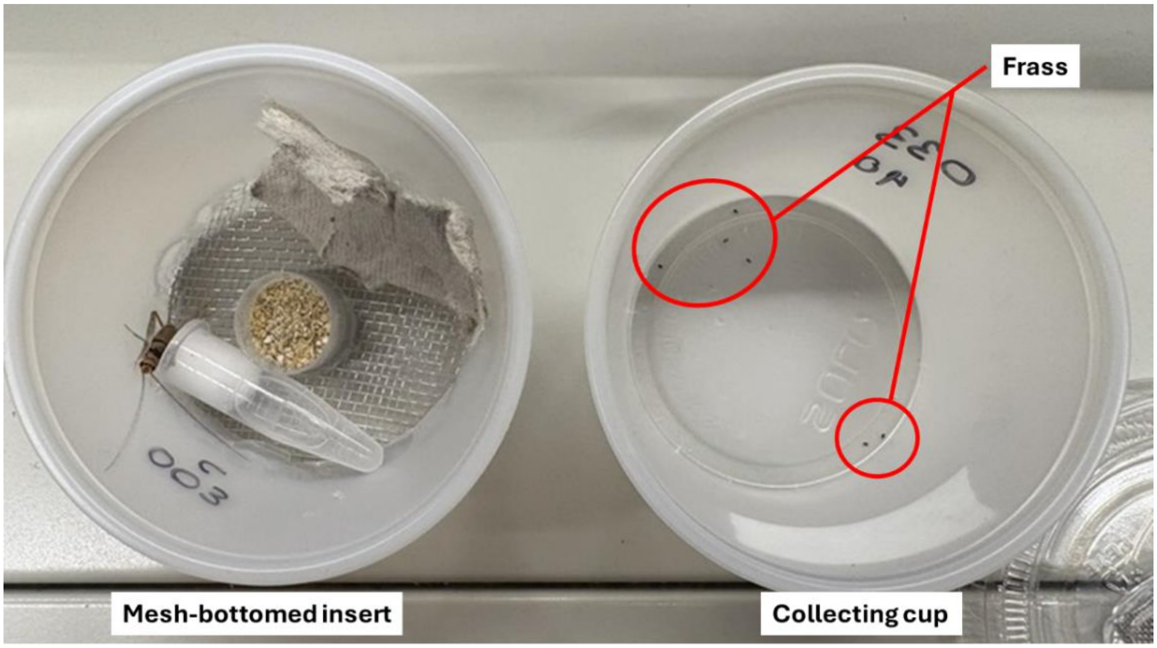


**Figure S3.** Individual sifting cup design. Crickets were housed in the mesh-bottomed insert cup (left) with shelter, food and water. The mesh-bottomed insert was nested within the collecting cup (right), where frass accumulated after dropping through the wire mesh.

**Table S1.** Type III ANOVA test for main effects and interactions in the mixed-effects cosinor model. Significant results for cos hour and sin hour indicate strong rhythmicity in amplitude and phase, while the significant diet × sin hour interaction indicates differences in phase timing among diets.

| Effect | $\chi^2$ | df | p-value |
|---|---|---|---|
| **Diet** | 1.15 | 3 | 0.765 |
| **Cos hour** | 96.16 | 1 | **<0.0001** |
| **Sin hour** | 129.19 | 1 | **<0.0001** |
| **Diet × cos hour** | 8.84 | 3 | **0.0314** |
| **Diet × sin hour** | 10.29 | 3 | **0.0162** |

**Table S2.** A linear mixed-effects cosinor model testing the effects of diet and rhythmic components on frass production. Estimates, standard errors (SE), degrees of freedom (df), t statistics, and p-values were obtained using Satterthwaite's approximation.

| Term | Estimate | SE | df | t-value | p-value |
|---|---|---|---|---|---|
| **Diet: MP** | –0.0219 | 0.1063 | 54.35 | –0.206 | 0.8377 |
| **Diet: Control - MP** | 0.0522 | 0.1063 | 54.25 | 0.491 | 0.6255 |
| **Diet: MP - Control** | 0.0787 | 0.1063 | 54.24 | 0.740 | 0.4623 |
| **Cos hour** | –0.4595 | 0.0469 | 1145.04 | –9.806 | **<0.0001** |
| **Sin hour** | 0.5179 | 0.0456 | 1137.39 | 11.366 | **<0.0001** |
| **Diet: MP × cos hour** | 0.0329 | 0.0653 | 1140.10 | 0.504 | 0.6142 |
| **Diet: Control - MP × cos hour** | –0.0997 | 0.0651 | 1139.63 | –1.530 | 0.1262 |
| **Diet: MP - Control × cos hour** | –0.1304 | 0.0651 | 1139.49 | –2.004 | **0.0453** |
| **Diet: MP × sin hour** | 0.1050 | 0.0636 | 1134.09 | 1.652 | 0.0989 |
| **Diet: Control - MP × sin hour** | 0.1599 | 0.0634 | 1134.02 | 2.522 | **0.0118** |
| **Diet: MP - Control × sin hour** | 0.1893 | 0.0634 | 1133.92 | 2.984 | **0.0029** |

### Cosinor model

Statistical equations and descriptive calculations were adapted from standard cosinor methodology as described in Hou et al. 2021 and Refinetti et al. 2007. Calculations for curve parameters for Figure 4 are as follows. Rhythmicity in frass production was quantified using a mixed-effects cosinor model. Frass mass at time $t$, $Y(t)$, was modeled as:

$$Y(t) = M_d + \beta_{\cos,d}\cos(\omega t) + \beta_{\sin,d}\sin(\omega t) + u_i$$

Where $M_d$is the MESOR (rhythm-adjusted mean) for diet $d$, $\beta_{\cos,d}$and $\beta_{\sin,d}$are the diet-specific cosine and sine coefficients, $u_i$is a random intercept for individual nymph identity, and $\omega = 2\pi/124$is the angular frequency of the 124-hour rhythm period determined from nonlinear least-squares estimation using **nlsLM** (minpack.lm). Diet-specific coefficients were estimated using a linear mixed-effects model:

$$Y(t) \sim \text{diet} \times (\cos(\omega t) + \sin(\omega t)) + (1 \mid \text{id})$$

Implemented using **lmer** (lmerTest). For the Control diet, $\beta_{\cos}$and $\beta_{\sin}$correspond to the model's main cosine and sine terms. For all other diets, coefficients were computed as:

$$\beta_{\cos,d} = \beta_{\cos} + \beta_{\cos\times d}, \beta_{\sin,d} = \beta_{\sin} + \beta_{\sin\times d}$$

Amplitude and phase were derived from the harmonic coefficients using standard cosinor transformations:

$$A_d = \sqrt{\beta_{\cos,d}^2 + \beta_{\sin,d}^2}$$

$$\phi_d = \text{atan2}(-\beta_{\sin,d}, \beta_{\cos,d})$$

Phase was optionally expressed in degrees as:

$$\phi_{d,\text{deg}} = \phi_d \times \frac{180}{\pi}$$

The timing of rhythmic extrema was computed from the phase. Peak time for each diet was obtained using:

$$t\_peak, d = (-\frac{\phi d}{\omega})\ mod\ 124$$

and trough time was defined as occurring half a period later:

$$t_{\text{trough},d} = \left(t_{\text{peak},d} + 62\right) mod\,124$$

Predicted frass values at these extrema were derived from the cosine waveform:

$$Y_{\text{peak},d} = M_d + A_d, Y_{\text{trough},d} = M_d - A_d$$

To provide a more interpretable measure of timing differences among diets, phase delay was computed as the difference in peak time between each diet and the Control diet:

$$\Delta t_{\text{peak},d} = t_{\text{peak},d} - t_{\text{peak,Control}}$$

Together, these cosinor-derived parameters quantify differences among diets in mean frass output (MESOR), rhythmic strength (amplitude), timing of rhythmic extrema (peak and trough times), peak–trough range, and waveform shape (β-coefficients, phase delay, rise–fall ratio), matching all values reported in Table S3.

**Table S3.** Descriptive properties of frass production across all diets summarizing observed frass behaviour during the fifth instar, including minimum and maximum frass mass (mg) and the duration of zero-frass periods within each diet. These values describe the basic distribution and zero-frass intervals in each diet group.

| Diet | Min frass (mg) | Max frass (mg) | Zero-frass period (h, min) | Zero-frass period (h, max) |
|---|---|---|---|---|
| Control | 0 | 4.1 | 12 | 36 |
| MP | 0 | 3.5 | 6 | 42 |
| Control–MP | 0 | 4.0 | 12 | 36 |
| MP–Control | 0 | 3.5 | 18 | 36 |

Note: MP had only one individual at 6 h and 42 h.

Descriptive measures of frass production during instar 5 showed that all diets produced similar minimum and maximum frass masses (0–4 mg) and comparable durations of zero-frass periods, although the MP group exhibited the widest range (6–42 h; Table S3), as evidenced by the gradual decline in frass output approaching ecdysis. These patterns indicate that total frass output was broadly consistent across treatments and time. Consistent with our statistical results, MESOR values were similar among diets (0.75–0.85 mg; Table 2B), reflecting the nonsignificant main diet effect on mean frass production. In contrast, the cosinor-derived parameters showed diet-related differences in both the magnitude of the rhythm (amplitude and peak–trough range, determined by $\beta\_cos$ and $\beta\_sin$ together) and its timing (phase, peak time, and phase delay, primarily governed by $\beta\_sin$), indicating that diets altered the structure of the frass production oscillation. Amplitude and peak–trough range were highest in both switched diets (amplitude: 0.88–0.92 mg; range: 1.75–1.84 mg), indicating stronger oscillations relative to the consistent diets. Diets also differed in phase timing, MP reached its peak approximately 2.47 hours earlier than Control, while the crossover diets were approximately 0.6–0.7 hours earlier. These timing shifts closely match the significant diet × sin interaction in the mixed-effects model

(Supplemental Table 1). Additional differences in waveform shape were reflected in the diet-specific β_cos and β_sin coefficients. Overall, both descriptive and rhythmic metrics demonstrate that while overall frass production was similar across diets, diet significantly influenced the strength, timing, and shape of the underlying frass rhythm.
